# Dual-modality imaging of immunofluorescence and imaging mass cytometry for whole slide imaging with accurate single-cell segmentation

**DOI:** 10.1101/2023.02.23.529718

**Authors:** Eun Na Kim, Phyllis Zixuan Chen, Dario Bressan, Monika Tripathi, Ahmad Miremadi, Massimiliano di Pietro, Lisa M Coussens, Gregory J Hannon, Rebecca C Fitzgerald, Lizhe Zhuang, Young Hwan Chang

## Abstract

Imaging mass cytometry (IMC) is a powerful multiplexed tissue imaging technology that allows simultaneous detection of more than 30 makers on a single slide. It has been increasingly used for singlecell-based spatial phenotyping in a wide range of samples. However, it only acquires a small, rectangle field of view (FOV) with a low image resolution that hinders downstream analysis. Here, we reported a highly practical dual-modality imaging method that combines high-resolution immunofluorescence (IF) and high-dimensional IMC on the same tissue slide. Our computational pipeline uses the whole slide image (WSI) of IF as a spatial reference and integrates small FOVs IMC into a WSI of IMC. The high-resolution IF images enable accurate single-cell segmentation to extract robust high-dimensional IMC features for downstream analysis. We applied this method in esophageal adenocarcinoma of different stages, identified the single-cell pathology landscape via reconstruction of WSI IMC images, and demonstrated the advantage of the dual-modality imaging strategy.

**Motivation:** Highly multiplexed tissue imaging allows visualization of the spatially resolved expression of multiple proteins at the single-cell level. Although imaging mass cytometry (IMC) using metal isotope-conjugated antibodies has a significant advantage of low background signal and absence of autofluorescence or batch effect, it has a low resolution that hampers accurate cell segmentation and results in inaccurate feature extraction. In addition, IMC only acquires mm^2^-sized rectangle regions, which limits its application and efficiency when studying larger clinical samples with non-rectangle shapes. To maximize the research output of IMC, we developed the dual-modality imaging method based on a highly practical and technical improvement requiring no extra specialized equipment or agents and proposed a comprehensive computational pipeline that combines IF and IMC. The proposed method greatly improves the accuracy of cell segmentation and downstream analysis and is able to obtain whole slide image IMC to capture the comprehensive cellular landscape of large tissue sections.

## INTRODUCTION

Understanding the cellular composition and its spatial distribution in tissue sections, termed “spatial biology,” is becoming increasingly important in a wide range of biological research fields^1^. Spatial biology utilizes highly multiplexed tissue imaging (MTI) techniques that allow visualization and quantification of the spatially resolved expression of various protein markers at single-cell resolution in tissues by acquiring multiple images of the same tissue that correspond to different markers, followed by computational deconvolution of the image data into single-cell-based marker quantification. MTI enables detailed spatial characterization of individual cells along with the complex interaction between target cells and their microenvironments, such as cancer cells and their neighboring immune and stromal cells. However, every MTI has platform-specific limitations that pose a substantial computational challenge to overcome these limitations; specifically, critical parameters in MTI are the number of multiplexed biomarkers and spatial resolution, which are mainly determined by the platform hardware.

Optical microscope-based platforms such as cyclic immunofluorescence (t-CyCIF)^2^, multiplex immunofluorescence (MxIF)^3^, multiplexed immunohistochemistry (mIHC)^4^, and CODetection by indEXing (CODEX)^5^ are optical microscope-based methods that usually have a high imaging resolution (0.05 to 0.25 μm^2^ per pixel). However, all these platforms achieve multiplex, ie. detection of multiple markers, via repeating the cycles of ‘antibody incubation’ – ‘imaging’ – ‘antibody removal’ in a sequential iterative process that usually takes 6 to 8 hours per cycle with repeated heating and cooling of the slide. The process, therefore, is prone to artifacts including autofluorescence (if detection is fluorescence-based), micro-shrinkage, expansion or loss of the tissue, and batch effect, which are exacerbated by the increasing number of cycles and markers. It becomes a critical issue especially when processing larger tissues (on the scale of cm^2^) as the micro-shrinkage and expansion are becoming non-negligible. It results in grave difficulties in downstream image registration and introduces significant errors. Moreover, autofluorescence and experimental batch effects also introduce biases into the data and may mask the true underlying biological signals^6–10^.

In contrast to the optical microscope-based MTI methods, mass spectrometry-based platforms such as imaging mass cytometry (IMC)^11^ and multiplexed ion beam imaging (MIBI)^12^ use metal isotope-conjugated antibodies and detect all markers in one round and raw images of all markers are acquired simultaneously. It is therefore free of the aforementioned artifacts that are caused by the repeated sequential cycles. Furthermore, these platforms have a significant advantage of low background signal, no autofluorescence, and no batch effect as the marker detection is not based on fluorescence intensity but time-of-flight (TOF) of the evaporating metal isotopes when the tissue is being ablated^11^. However, IMC has a much lower spatial resolution (1 μm^2^ per pixel) that hampers accurate cell segmentation, resulting in inaccurate single-cell feature extraction and hindering accurate cell phenotyping^13,14^. Baars and colleagues published a method that used high-resolution IF as a segmentation called MATISSE (iMaging mAss cytometry mIcroscopy Single cell SegmEntation)^15^. However, this method is still restricted by the confined IMC field of view (FOV), as the platform hardware only acquires mm^2^-sized rectangular regions. For larger cm^2^-sized tissue, the acquisition needs to be broken into a number of smaller separate rectangular FOVs, which greatly limits its application in studying large whole-tissue cellular landscapes, such as an entire patient tumor or an intact mouse cerebrum.

Here, we developed a comprehensive solution that not only significantly improved the accuracy of cell segmentation but also was able to reconstruct multiple separately acquired IMC FOVs to a single WSI so the tissue’s cellular composition and spatial landscape could be studied as a whole. We demonstrated the method using endoscopic mucosal resection (EMR) specimens that contained esophageal adenocarcinoma and adjacent precancerous Barrett’s esophagus. The results faithfully recapitulated the pathological transition of the cellular landscape between Barrett’s esophagus, dysplasia, and adenocarcinoma that was only possible to study as a whole in WSI.

## RESULTS

Our method aimed to combine the advantages of IF and IMC by dual-modality imaging of the same tissue, and then reconstruct the small rectangle IMC FOVs to WSI based on the IF reference to analyze the whole tissue with both high-dimensional and high-resolution imaging data (Figure 1A). To demonstrate its applicability in different tissue types, we stained large endoscopic mucosal resection (EMR) tissues of early esophageal adenocarcinoma with adjacent precancerous Barrett’s esophagus, which represents various histologic features, including squamous and glandular epithelia and stromal tissue, and pathological progress from normal tissues to metaplastic precancerous lesion, dysplasia, and cancer in a large area measuring approximately 0.5 cm x 2 cm (Supplementary Figure S1).

**Figure 1.**
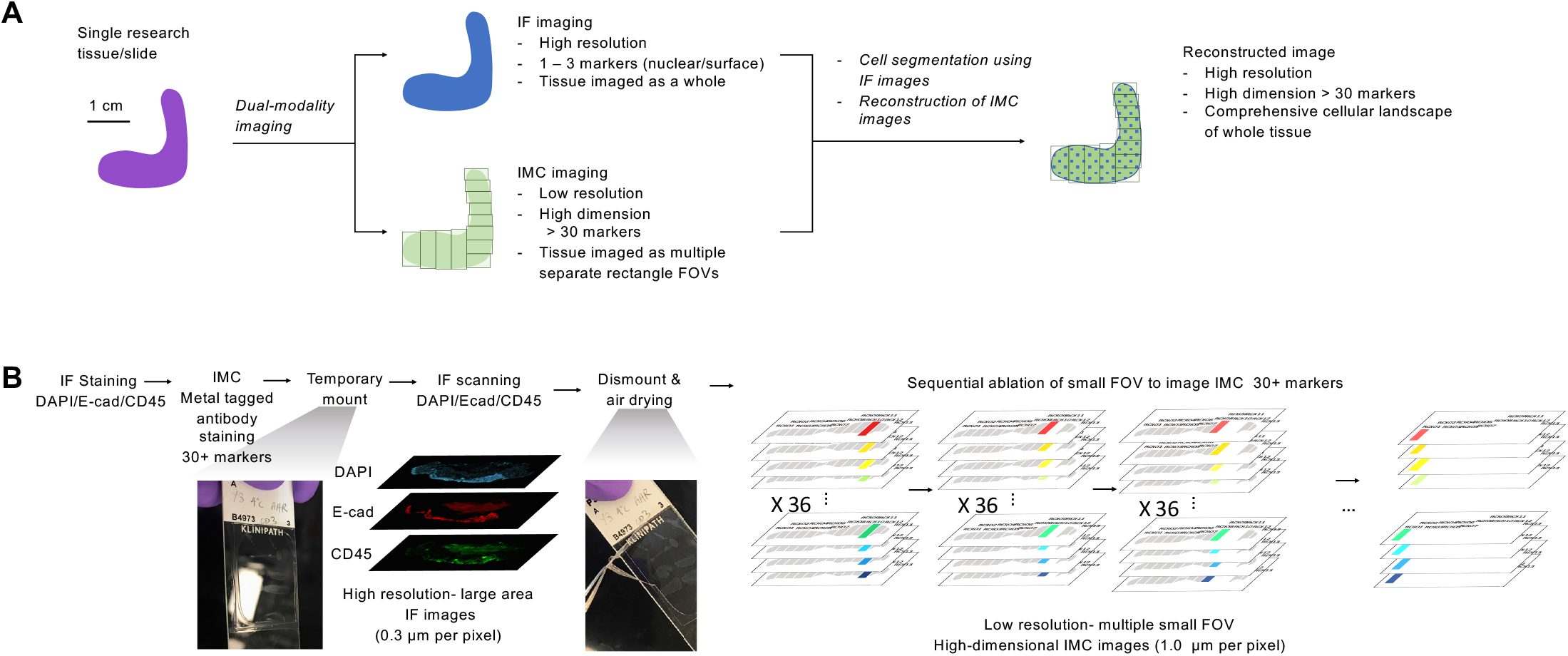
Experimental benchwork workflow for integrating dual-modality imaging. A. The method principle: the same tissue sample was imaged for both IF and IMC. IF images provided a high-resolution image for accurate cell segmentation and whole tissue reference, yet only for up to three markers; IMC images provided a high-dimensional image dataset for complex spatial cell phenotyping but were acquired as separated small rectangle FOVs that depends on the tissue shape and size. We combined the advantages of the two imaging modalities with accurate cell segmentation and reconstruction of high-dimensional image data for whole tissue. B. Experimental procedure: the tissue slide was stained first for IF E-Cadherin and CD45, followed by staining of 36 metal-tagged IMC antibodies. Then, a temporary mounting was performed to scan the IF images. IF optical scanning was performed to obtain high-resolution (0.3 μm per pixel), whole-area IF images. After dismounting and air drying, IMC markers were acquired as separate small rectangle FOVs (1.0 μm per pixel). The two datasets were then processed for cell segmentation and FOV stitching.

### Staining and dual-modality imaging of IF and IMC

Briefly, we stained the slide for two rounds: first with the primary IF antibodies (E-Cadherin, CD45), and second with the secondary IF antibodies and a cocktail of metal isotope-conjugated IMC antibodies (36 markers, Supplementary Table S1). The slides were stained with DAPI before temporarily being mounted for whole slide fluorescent scanning for IF markers, followed by dismount and air drying, and acquisition of multiple small rectangle FOVs of IMC (Figure 1B and **METHOD**). The hands-on time for the benchwork was approximately 2 hours, and the full preparation of the slide could be finished in one day.

### Computational integration pipeline of IF and IMC images

It has been difficult to digitally stitch separately acquired FOV IMC images even if they were adjacent because, in IMC, imaged areas are destroyed during acquisition therefore there are no overlapping areas that could be used as references. We developed a computational approach to map mm^2^-sized smaller FOV IMC images onto cm^2^-sized larger WSI IF images, and then register and stitch them together (Figure 2A). Briefly, we first determined the global coordinates of small FOV IMC images by identifying high pixel-level correlation with the reference IF WSI and then refined the registration locally between small FOV IMC and IF. By using our registration method (see **METHOD**), we successfully registered high-dimensional IMC images with 36 markers (Supplementary Figure S2) onto WSI IF with a high subcellular resolution (Figure 2B).

**Figure 2.**
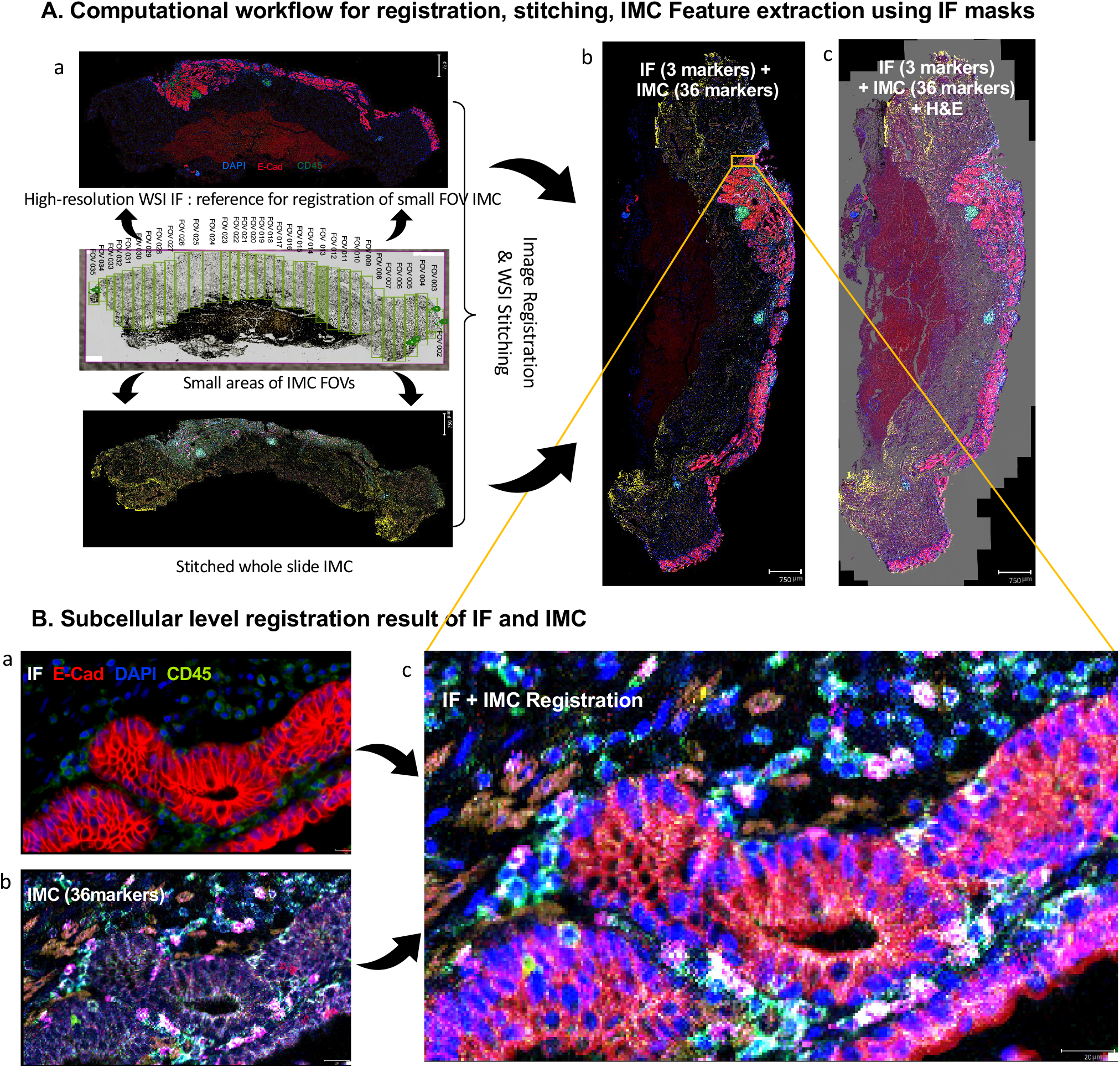
Image registration and stitching of IMC to IF. A. Computational workflow for registration and stitching of IMC to IF images (a) Whole slide IMC images are generated by stitching and registering small FOV IMC images to large-sized IF images as a reference. (b) 3 IF markers and 36 IMC markers are visualized on one slide by registration. (c) Registration of H&E images to the IF and IMC B. Registration of IF (DAPI, E cadherin, CD45) and IMC images (36 antibodies) at subcellular levels. (a) IF images of E-Cadherin, DAPI, and CD45. (b) IMC images of 36 markers. (c) Subcellular registration of IF and IMC images

### Generation of customizable region of interest (ROI) masks from WSI IMC images

Once the whole tissue was reconstructed for both IF and IMC, we further registered the image with an adjacent slide stained for hematoxylin and eosin (H&E). The registered H&E image is important for clinical samples that pathologists, who rely on H&E images to make the final diagnosis, are able to define a disease-specific region. For example, different pathological stages with any shapes (Figure 3A). These areas could then be extracted from the WSI IMC images and individually studied for their cellular composition (Figure 3B).

**Figure 3.**
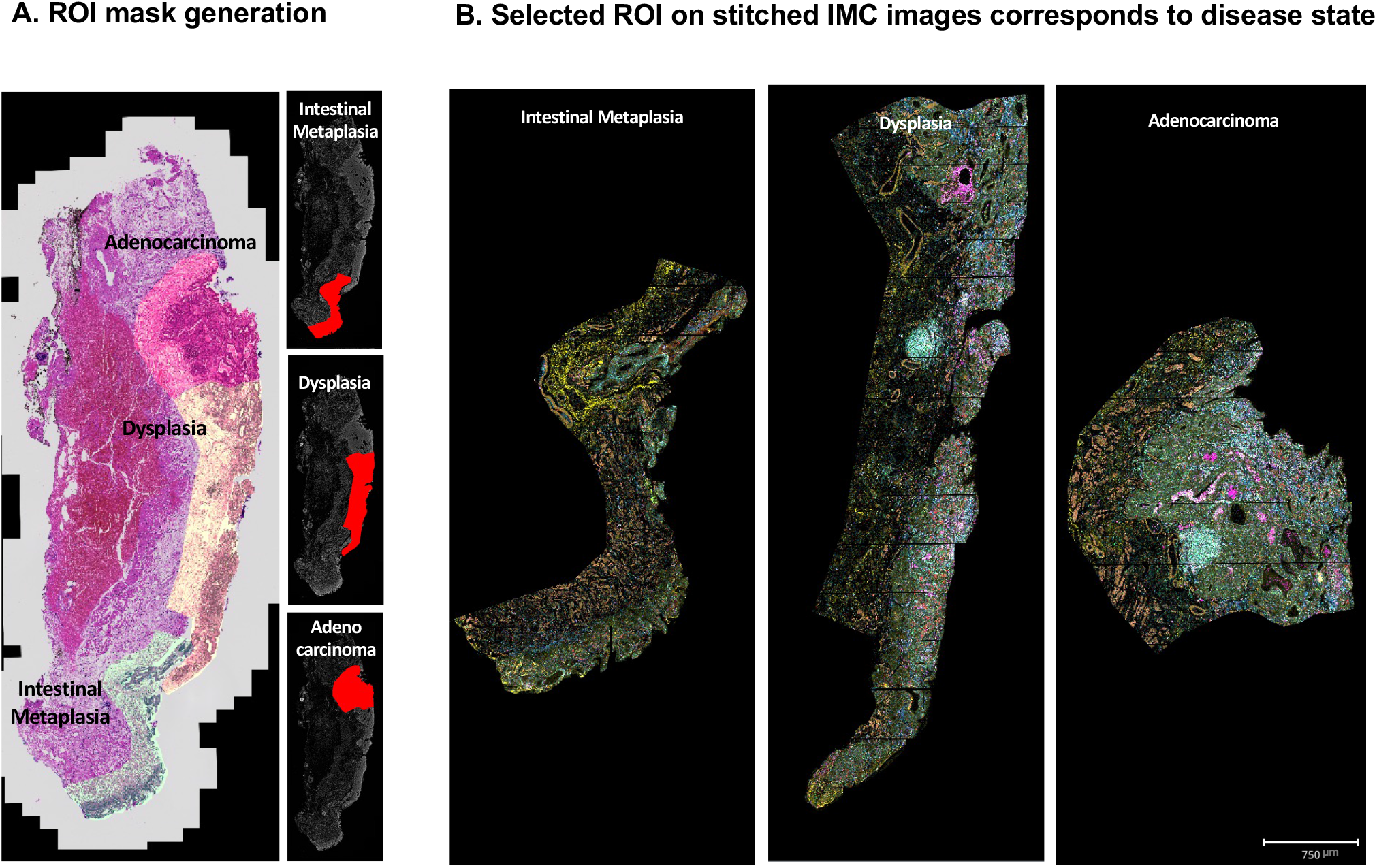
Generation of the ROI Mask according to the disease status from whole slide IMC. A. Generation of a new disease status mask from the co-registered H&E slide. B. Application of the disease status mask to IMC images and reconstruction of IMC images according to the disease sequence.

### Generation of accurate cell masks using registered IF

In order to establish accurate cell segmentation for low-resolution IMC images, we generated cell masks using MESMER, a deep learning-based algorithm for nuclear and whole-cell segmentation^16^, from the registered high-resolution IF using IF DAPI for the nuclear marker and IF E-Cadherin for the epithelial cell membrane marker. We registered IF-based cell segmentation masks with IMC images so that we could extract high-dimensional, autofluorescence-free IMC marker intensity features from accurate cell segmentation boundaries for downstream analysis (Figure 4).

**Figure 4.**
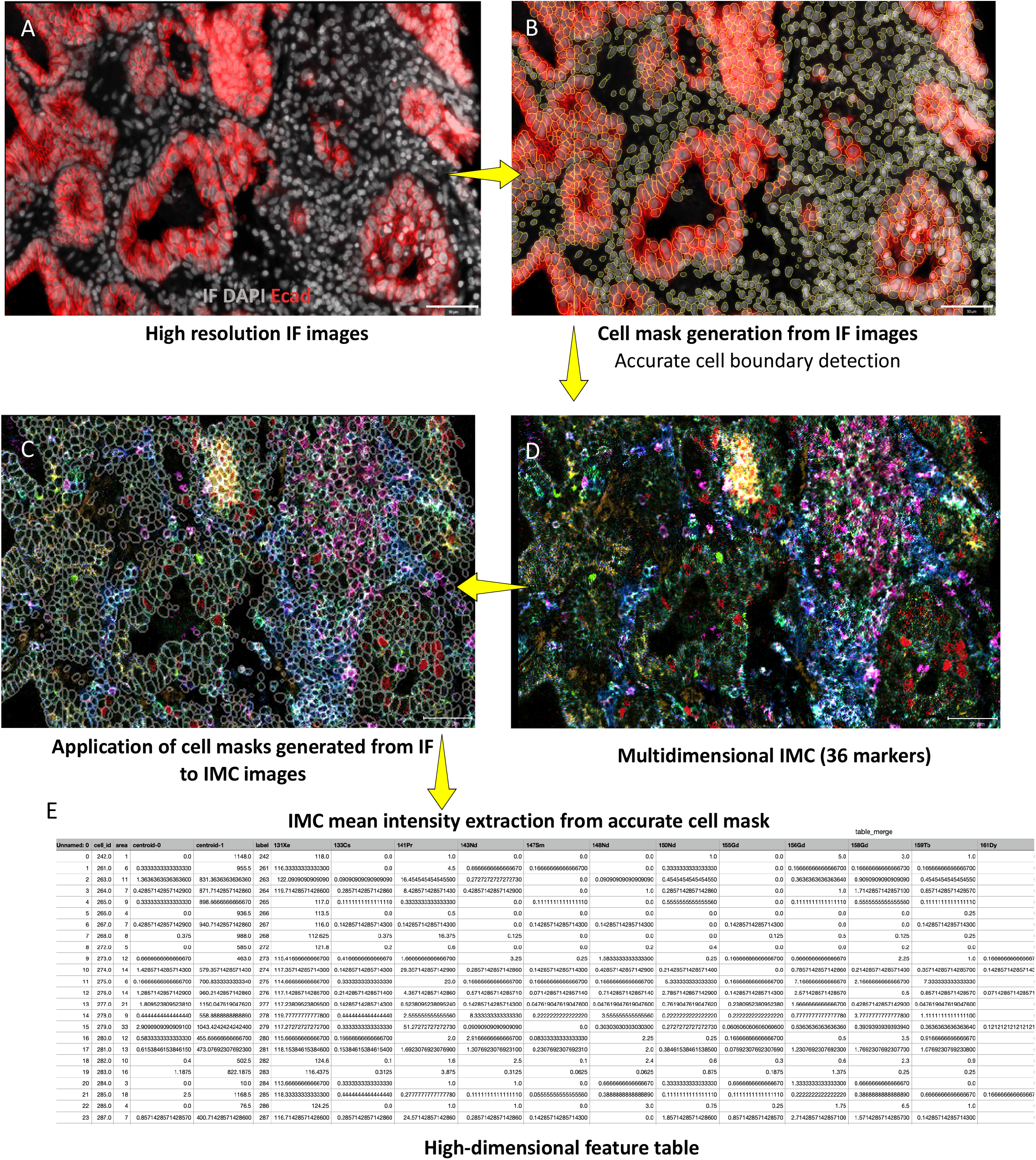
Generation of accurate cell masks using registered IF. (a) High-resolution IF images (IF; DAPI, and E Cadherin). (b) Generation of accurate cell segmentation masks from high-resolution IF images. (c) Multidimensional IMC images of 36 markers. (d) Application of cell masks generated from IF to IMC images. (e) Application of cell masks to high-dimensional IMC images for obtaining a high-dimensional feature table at the single cell level.

We then compared segmentation masks generated from high-resolution IF versus low-resolution IMC and found the IF mask resulted in highly accurate demarcation of the true boundaries for both cell and nucleus across all tissue types and disease stages (Figure 5A and Supplementary Figure S3). In addition, cytoplasmic areas were more clearly delineated in IF-based cell segmentation than in IMC-based (Figure 5A).

**Figure 5.**
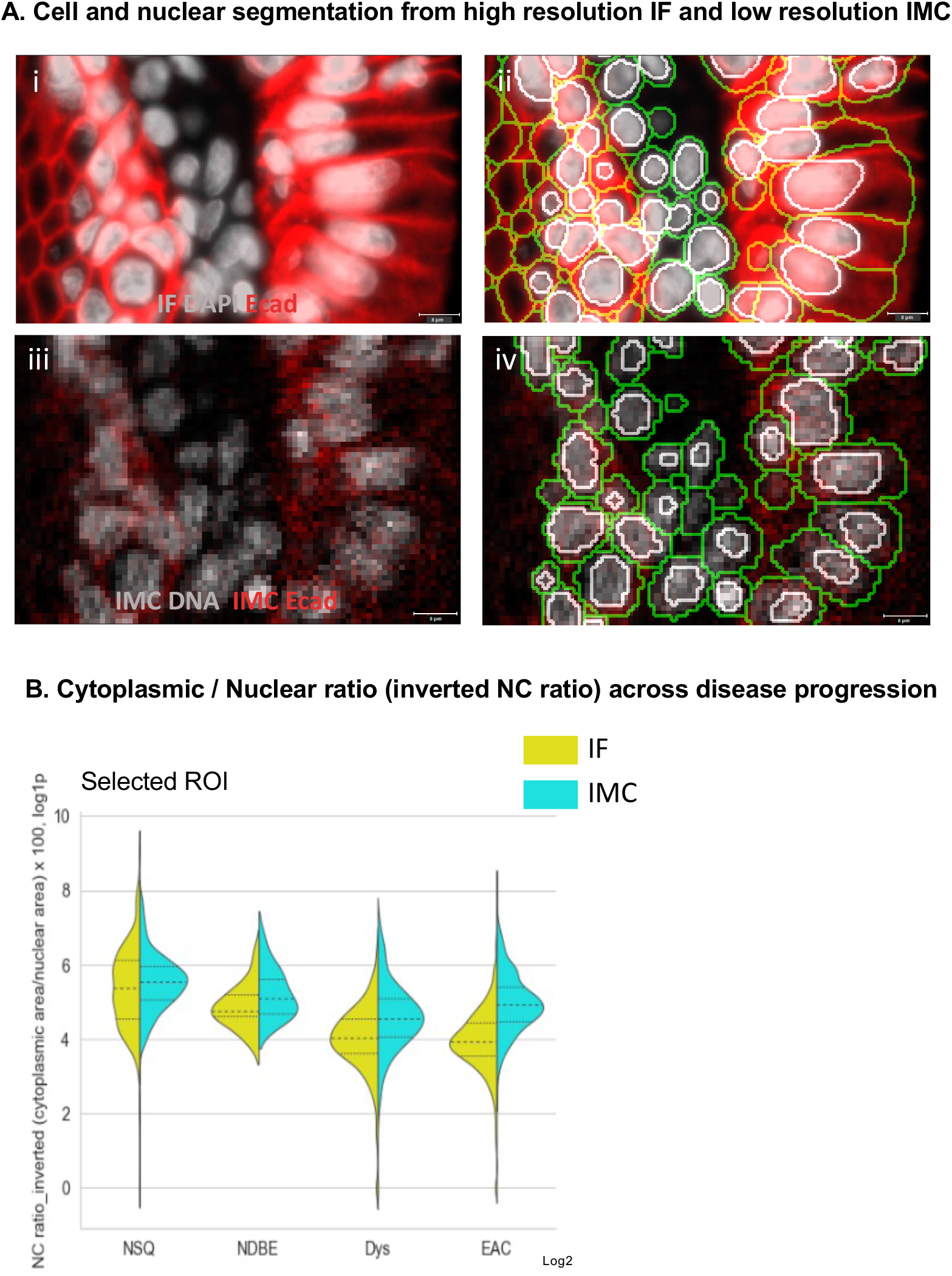
Comparison of the cell and nuclear masks between IF and IMC images. A. Cell and nuclear segmentation from high-resolution IF and low-resolution IMC. (i) High-resolution IF images (gray, IF DAPI, red, E cadherin). (ii) Cell (green) and nuclear (white) masks from high-resolution IF images. (iii) Low-resolution IMC images (gray, IMC DNA, red, IMC E-Cadherin). (iv) Cell (green) and nuclear (white) masks from low-resolution IMC images. B. Cytoplasmic/Nuclear ratio (inverted N/C ratio) across disease progression in selected regions of interest.

We further compared the accuracy of IF-based and IMC-based segmentation masks by evaluating the inverted nucleus-to-cytoplasmic (N/C) ratio in different disease statuses. Histomorphologically, the inverted N/C ratio decreases along with disease progression from intestinal metaplasia to dysplasia and adenocarcinoma^17^, which was faithfully recapitulated by the IF-based cell segmentation but not IMC-based (Figure 5B), indicating that IF-based segmentation is sufficiently sensitive for subcellular phenotyping, especially for clinical samples.

The robustness of cell segmentation was further evaluated by biaxial plots of mutually exclusive marker pairs, such as PanCK (epithelial cells) and αSMA (vascular, fibroblast, smooth muscle cells), whereby double positivity indicates marker spillage that is usually caused by imperfect segmentation. Interestingly, both IF-based and IMC-based cell segmentation showed very low double positivity on two mutually exclusive marker pairs, PanCK/αSMA and PanCK/CD4 (Supplementary Figure S5). However, IF-based segmentation was able to detect 199,425 cells from the tested region while the IMC-based detected 120,543 cells, which generally suggested that IF-based segmentation was more sensitive to distinguish individual cells that were particularly important when studying tissues with densely packed cells, such as tumor.

Finally, we evaluated the batch effect in IF and IMC by assessing the signals of the same markers that were acquired on different experiment dates. While each experiment was strictly controlled that staining and imaging were performed with the same parameters, we found the batch effect was inevitable for IF markers that varied intensities were observed in different batches (Figure 6A). On the other hand, the signal intensity of IMC markers was highly consistent and free of batch effect (Figure 6B), indicating IMC image data, even acquired in different batches, could be analyzed together with little need for batch correction, which highlights the advantage of IMC for studies with larger sample cohorts that image data need to be acquired in multiple batches.

**Figure 6.**
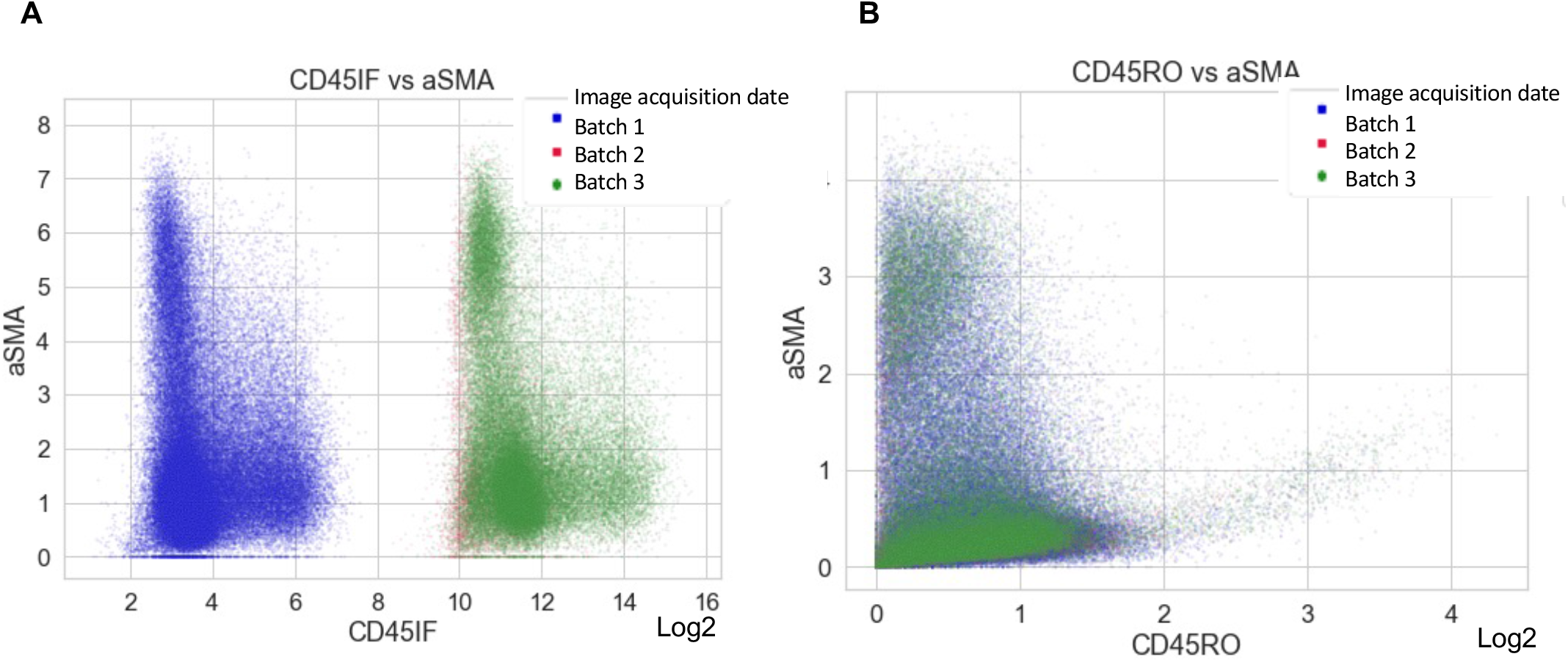
Batch effect of IF vs. IMC. A. Scatter plot of mean intensity between IF CD45 and IF αSMA according to the image acquisition date. B. Scatter plot of mean intensity between IMC CD45RO and IMC αSMA according to the image acquisition date. The batch effect was found only in IF (A) and not in IMC (B).

### Accurate cell phenotyping results based on IF-based cell mask

We compared IF-based versus IMC-based segmentation in downstream analysis. Briefly, single cells extracted from the same high-dimensional image dataset using either IF-based or IMC-based segmentation were pooled together and analyzed for cell phenotyping using the GPU-boosted implementation of PhenoGraph (unsupervised clustering approach)^7^. All the cells were grouped into 19 clusters. Interestingly, IF-based segmentation was able to detect a higher number of Cluster 5 (highlighted in Figure 7A). Cluster 5 showed high PanCK, and moderate CD1a and CCR7 expression. To further confirm the cell identity, we visualized the in situ distribution of cluster 5 using the open-source visualization tool Napari (https://github.com/napari/napari) (Figure 7B) and found the cells are enriched in the suprabasal layers in the squamous epithelium of the normal esophagus, which were likely to be the Langerhans cells or a Langerhans cell subset^18^. It was clear that the high-resolution IF-based segmentation provided a more accurate cellular basis for high-dimensional downstream phenotyping that the details of cell location and morphology were maintained.

**Figure 7.**
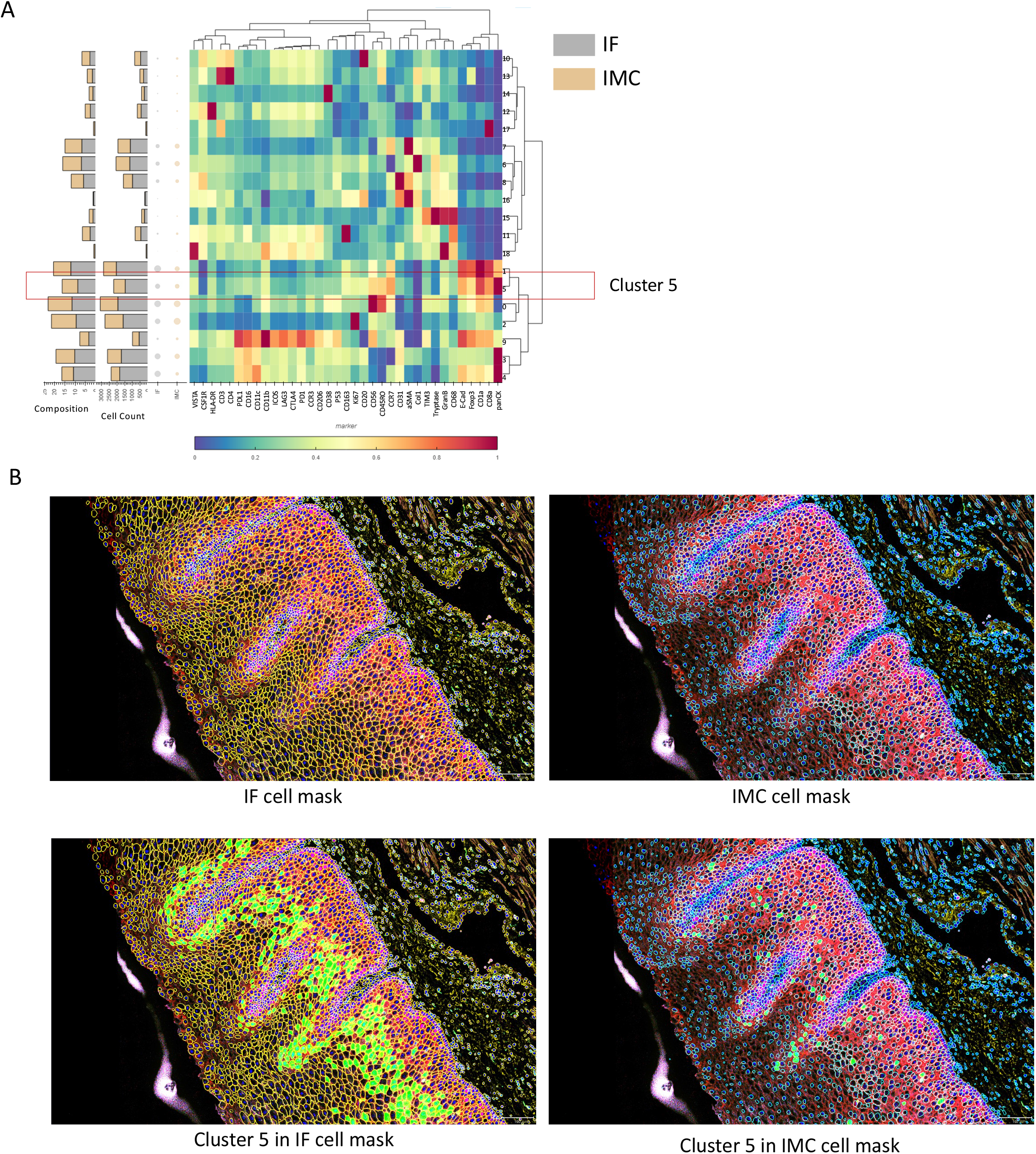
Comparison of the downstream analysis of unsupervised clustering results between IF-based cell mask and IMC-based cell mask. A. Unsupervised clustering using GPU-boosted implementation of PhenoGraph from the extracted mean intensity from the IF mask (gray) and the IMC mask (light brown). B. Mapping of cluster 5 on the IF cell mask and the IMC cell mask.

## DISCUSSION

### Summary of this study

In this study, we present a highly practical approach that integrates mass spectrometry-based IMC with optical microscope-based IF that exploits the technical advantages of both imaging platforms while overcoming their own limitations. Our method is capable of analyzing tissues of varying sizes and shapes with highly accurate cell segmentation (based on IF images) and high dimensional and large-scale phenotyping (> 30 markers based on WSI IMC images) without introducing autofluorescence or batch effects. By stitching smaller FOVs to obtain a larger WSI, our method overcomes the main limitation in IMC which only acquires small rectangular FOVs with low resolution.

### A practical solution of integrating IF into the IMC imaging system

Our staining protocol enables dual staining of IMC and IF on the same slide, which was enabled by a relatively simple tweak of the mounting method; also, our computational integration tool was designed based on a widely used open-access computational platform, which is easy and convenient for IMC users to employ our tool in their analysis. In addition, compared with metal-conjugated IMC antibodies, IF antibodies are highly available and widely validated. In our system, key markers could be detected by IF antibodies if the IMC antibodies are not available. Also, user-conjugated IMC antibodies with a new metal tag can be validated using the corresponding IF staining on the same slide, which facilitates the development of new antibody panels in IMC.

### Advantages of large-size reconstructed IMC images

The registered and stitched WSI IMC images with adjacent H&E enable pathologists to examine and understand the semantic, structural, and spatial context with regard to the sequential progression of diseases, especially the subtle and transitional changes that appear across large areas. WSI IMC images also allow the selection of representative and customizable ROIs of the researcher’s interest, instead of using small and rectangle FOV of IMC images. The shape of an ROI, such as a concave shape, can significantly affect spatial analysis, including neighborhood enrichment analysis and shortest average distance. It is therefore important to have customizable area masks that avoid this bias. In addition, the cell population composition or cell number can be accurately compared for a meticulously drawn ROI via pathologic semantic evaluation from large WSI-level IMC, thus enabling robust comparison between different tumor microenvironments.

### Advantage of the integration of IF and IMC modality in terms of imaging processing time

Our new integration method is time-efficient in acquiring image data for WSI. In optical microscope-based platforms such as t-CyCIF^2^, the maximal number of antibodies for one cycle is usually 3-5 markers, and one channel should be nuclei staining such as DAPI as it is required for image registration. Therefore, for staining a large tissue with 36 markers, 9 rounds with 4 antibodies each are needed. Moreover, as a stripping-out process is needed between scanning and new staining during the cycles, it takes multiple days for image acquisition^19,20^.

In this regard, IMC has a significant advantage over cyclic multiplexing imaging in terms of image acquisition time and automation. Unlike the cyclic imaging modality, IMC is an all-in-one staining and all-in-one imaging method using a single master mix of metal-conjugated primary antibodies and does not require repeated sequential steps such as antibody staining, image acquisition, and antibody removal^21^. A complete set of antibodies with more than 50 markers can be applied simultaneously, and there is no need for manual intervention during the acquisition. In addition, there is no tissue loss or micro-expansion issue from repeated antibody removal steps. As the hands-on time for the benchwork is approximately 2 hours, acquiring multiple FOVs for WSI imaging can be completed within one day (time for acquiring 0.5 mm x 0.5 mm area at 1-μm resolution: ~3.5 hours); in our case, about 1.5 days were needed for the entire processing of IMC imaging for a whole slide with 36 antibody markers. In addition, unlike slides generated from fluorescence-based imaging modalities, slides stained with IMC can be stored for a long time without degradation.

### IF combination improved the accuracy of cell segmentation and downstream analysis

Although IMC has many advantages, IMC images have a relatively low resolution (cellular resolution: 1 μm/pixel^11^) compared to high-resolution optical-based images such as CyCIF or MxIF (cellular resolution: 0.325 μm/pixel). Low-resolution images hamper accurate cell segmentation, induce lateral spillage of cell surface expression from neighboring cells, and result in inaccurate single-cell feature extraction. As a result, mutually exclusive markers are often detected in one cell (i.e., false-double positive cells), which hinder accurate cell phenotyping^14^ and subsequent downstream analysis. To avoid this issue, a recent study (REDSEA) proposed a new method for correcting cell phenotyping by assigning the signal in the cell boundary to each cell. However, there are still many difficulties in detecting accurate cell boundaries with low-resolution IMC images^14^. In this regard, our results showed that IF-based cell segmentation lowered the rate of double-positive cells (Supplementary Figure 5).

Nuclear imaging of IF DAPI most effectively revealed the nuclear structure at high resolution, delineating nuclear contour and nucleus texture even in densely packed cells, intermixed immune cells, and membrane imaging of IF E-Cadherin delineated stratified normal squamous epithelium without a nucleus (Figure 5A). IF DAPI is considered the empirical ground truth as the most widely used marker in nuclear segmentation and image registration^9^. We were able to generate accurate nuclei and cell masks using high-resolution IF DAPI and E-Cadherin images and register them with IMC images, which enabled us to analyze high-dimensional IMC images using accurate IF-based cell masks.

### Comparison with other multiplexed methods using integrated imaging modalities

A few new spatial multiplexed technologies have integrated multiple platforms to improve research outcomes. NanoString Digital Spatial Profiling (DSP)^22^ uses sequencing instead of an imaging-based method to analyze RNA or protein (oligo-tagged antibodies). IF imaging is used to define specific ROIs, such as pan-cytokeratin stained epithelial area or CD45 stained immune cell area, from which multiplexed tags and/or oligos could be retrieved and sequenced, therefore the spatial profiling is achieved. It is noteworthy that the smallest spatial region NanoString DSP can analyze is 40 to 800 μm, therefore it does not achieve a single-cell level. On the other hand, image results from our method could be analyzed to the subcellular level thanks to the high-resolution IF. MATISSE (iMaging mAss cytometry mIcroscopy Single cell SegmEntation)^15^ uses a similar concept that integrated IF with IMC to increase image resolution for cell segmentation. However, it used only nuclear marker IF DAPI and was still restricted by the small rectangle FOV in IMC. In our method, DAPI and other IF markers could be included, such as lineage and/or membrane markers such as E-Cadherin and CD45 that greatly facilitate the downstream cell segmentation or specific markers of interest that are not available in IMC. It’s also noteworthy that our method allows the reconstruction of large whole tissues of high-resolution IF and high-dimensional IMC images with a registered H&E image. This is a great advantage in studying large clinical tissues that detailed pathological review is needed.

### Limitations of Study

There are limitations in this study that is also shared by many spatial multiplexed imaging technologies, whereby cells, which are three-dimensional, round, and voluminous, are cut with a microtome into two-dimensional planes for imaging analysis. It inevitably leads to variations in the observed morphology depending on which plane the cells are cut from. Therefore, there was a limitation in fully evaluating the cell morphology, such as the N/C ratio, even under high-resolution images. This may be resolved by further development of three-dimensional spatial imaging technologies.

## Acknowledgments

This work was supported by the CRUK-OHSU joint grant to YC and LZ (C65718/A29808). YC and LC acknowledge funding from the National Institutes of Health (1U01 CA224012, U2C CA233280), the Knight Cancer Institute, and the OHSU-Brenden-Colson Center for Pancreatic Care. YC also acknowledges funding from the National Institute of Health (U54CA209988, R01 CA253860). The laboratory of RF is funded by a Core Programme Grant from the Medical Research Council (RG84369). DB and GH are funded by the IMAXT Cancer Grand Challenge grant (A24042). GH is funded by a Royal Society Research Professorship (RSRP\R\200001), a Wellcome Trust delta-tissue Leap grant (G113035), and a Cancer Research UK Core funding through the University of Cambridge’s Cancer Research UK Cambridge Institute (C9545/A29580) A24042, A21143.

We thank all patients who contributed to the study. We thank Tara Evans, Michele Bianchi, Bincy Alias, and other members of the NIHR Cambridge Clinical Research Centre for their assistance in collecting the EMR tissues. We thank the Human Research Tissue Bank, which is supported by the UK National Institute for Health Research (NIHR) Cambridge Biomedical Research Centre, from Addenbrooke’s Hospital. We acknowledge the infrastructure support from the Experimental Cancer Medicine Centre and NIHR Cambridge Biomedical Research Centre (BRC-1215-20014).

## Author contributions

Eun Na Kim performed the data analysis and prepared the manuscript and figures.

Phyllis Zixuan Chen performed the IMC experiments and optimized the integration of IMC and IF methods. Monika Tripathi and Ahmad Miremadi assessed the EMR tissue and graded the local pathological grades. Massimiliano di Pietro supervised the collection of the endoscopic samples.

Lisa M Coussens and Rebecca C. Fitzgerald conceived the study design and analysis and provided guidance on preparing the manuscript.

Young Hwan Chang established the computational pipeline of registration.

Young Hwan Chang and Lizhe Zhuang conceived the study design and analysis, performed the analysis, supervised the research, and prepared the manuscript and figures.

## Declaration of interests

The authors declare no competing interests.

## STAR METHODS

### Ethics statement

The studies involving human participants were reviewed and approved by the East of England - Cambridge Central Research Ethics Committee. The patients/participants provided their written informed consent to participate in this study.

### Resource availability

The imaging data generated during this study are available at https://doi.org/10.5281/zenodo.7576005. The published article includes all datasets generated or analyzed during this study. Lead contact Further information on methodology and code should be requested and will be fulfilled by the lead contact, Young Hwan Chang.

### Materials availability

The original contributions presented in the study are included in the article/Supplementary Material. Further inquiries can be directed to the corresponding authors.

### Data and Code Availability

All software used in this manuscript is detailed in the article’s Methods section. Our code (in Matlab and Python) is available at https://github.com/enkim/IMC-IF.

## METHOD DETAILS

### 1. Tissue samples

We used endoscopic mucosal resection (EMR) samples from a pilot cohort of early EAC with Barrett’s metaplasia (n=3) and normal esophagus (n=2) from tissue donations. The EMR tissues contain sequential stages of tumorigenesis including normal esophageal squamous epithelium (NSQ), Barrett intestinal metaplasia (NDBE), low-and-high grade dysplasia (Dys), and esophageal adenocarcinoma (EAC, Supplementary Figure 1). The study was approved by the Institutional Ethics Committees and all subjects provided informed consent for the use of their tissue samples for research purposes (REC 01/149).

### 2. Pathology diagnosis for Barrett’s esophagus

Surgical pathologists independently assessed the pathology as described in our previous study^23^. The H&E slides were scanned and printed out. Two experienced gastrointestinal pathologists (M.T. and A.M.) independently assessed the H&E slides using the definitions and histological criteria for NSQ, NDBE, Dys, and EAC recommended by The Royal College of Pathologists^24^ and guidelines by the British Society of Gastroenterology on the diagnosis and management of Barrett’s esophagus^25^. High- and low-grade dysplasia were grouped together as dysplasia.

### 3. Antibody

We used three IF antibodies (CD45, E-Cadherin, Pan-cytokeratin) and 36 IMC antibodies targeting lineage markers (Pan-cytokeratin, E-Cadherin, Alpha-SMA, Col1, CD31), cell status markers (P53, Ki67), and immune cell phenotyping markers (CD45RO, CD3, CD4, CCR3, CCR7, CD56, CD8a, Granzyme B, ICOS, TIM3, LAG-3, CTLA4, FoxP3, PD-1, CD20, CD38, CD11b, CD16, Mast cell tryptase, CD11c, CD14, HLA-DR, CD68, CD163, Mannose Receptor, CD1a, CSF-1-R, PD-L1, VISTA, Supplementary Table 1).

### 4. Antibody validation

A full panel validation was performed^21^. All antibodies underwent extensive validation prior to multiplexing. A pathologist visually evaluated and validated the staining pattern of all IMC and IF antibodies by comparing the alleged positive and negative controls from the human protein atlas (https://www.proteinatlas.org). We assessed whether single-plex IHC using identical antibodies and 36-plex imaging mass cytometry analysis yielded the corresponding results^11^ between unlabeled and metal-labeled antibodies. Antibodies were individually validated to verify target specificity.

### 5. New mounting technique for dual-modality imaging

We developed a new staining/mounting technique that allows both IMC (36 markers) and IF (DAPI/CD45/E-Cadherin) in the same slide section without harming/destroying the tissue. The proposed approach consists of a benchwork for image data acquisition and a computational pipeline for image data integration. The benchwork (Figure 1A) is a practical tweak of the IF and IMC staining protocols in which the slide is stained for three rounds of antibodies in the order of primary cell surface marker antibodies (E-Cadherin in this study), secondary fluorescent antibodies (Alexa fluor 647 in this study) and a cocktail of metal isotope-conjugated IMC antibodies (36 markers in this study). The slides are temporarily mounted for whole-slide fluorescent scanning, followed by dismounting and air drying; the dried slides can be stored for months at 4°C. In addition, depending on the slide scanner, two more markers can usually be added for other cell types, such as CD45 for immune cells or CD31 for endothelial cells. The hands-on time for the benchwork is approximately 2 hours. For optimal results, we recommend incubating the primary antibodies at 4°C. However, the incubation may be reduced to 1 to 2 hours per round so that full preparation of the slide could be finished in one day.

a. IF staining. Dewax and antigen retrieval were performed on formalin-fixed paraffin-embedded (FFPE) EMR tissues section as described^23^. The slides were blocked by 5% BSA in PBS for 45 minutes at room temperature, followed by incubation of IF primary antibodies of CD45 (#13917, Cell Signaling Technology, 1:100) and E-Cadherin (#14472, Cell Signaling Technology, 1:100) that diluted in PBS-TB (PBS supplemented with 0.05% Tween 20 and .05% BSA) overnight at 4C. It was advised to prepare all buffers and reagents in plastic containers instead of glass bottles to avoid contamination of metal traces.
b. IMC antibody staining with 36 markers. The slides were washed in PBS for 3 x 5 minutes, followed by incubation of IF secondary antibodies for CD45 (Thermo Fisher Scientific, A21428, 1:200) and E-Cadherin (Thermo Fisher Scientific, A21240, 1:200) for 1 hour at room temperature. The slides were then washed for 3 x 5 minutes and stained for IMC antibody cocktail that was prepared in PBS-TB overnight at 4C.
c. Temporary mounting. The slides were washed for 3 x 5 minutes and incubated with Intercalator-Ir (Cell-ID™, 201192B, Standard BioTools Inc, 1:400) and DAPI (final concentration of 0.5 μg/mL) in PBS-TB for 5 minutes at room temperature. The slides were briefly washed once in PBS and mounted in 5% BSA in PBS. The edge of the coverslips was sealed with rubber cement (29010017000, Marabu Fixogum) and then air dried for 20 minutes.
d. IF image acquisition. The slides were then scanned in Zeiss Axio Scan Z1 for IF images for CD45, E-Cadherin, and DAPI. In this study, we chose a 20X objective for an image resolution of 0.325 μm per pixel. The image acquisition took approximately 45 minutes for one EMR tissue with a 0.5 x 1 cm size. 40x objective for a resolution of 0.163 μm per pixel was also available. The IF images were exported as TIFF format using ZenLite software.
e. Dismount. The dried rubber cement was stripped using forceps and the slides were submerged in PBS-T (PBS supplemented with 0.05% Tween 20) to dismount the coverslips. The slides were then washed in PBS-T for 3 minutes and then in Milli-Q water for 3 minutes. The slides were then air-dried for 15 minutes at room temperature and stored at 4C until IMC acquisition.
f. IMC data acquisition and processing (1 μm per pixel): Images were acquired using the Hyperion Imaging System (Standard BioTools Inc, previously Fluidigm). Each field of view (FOV), which was an adjacent plane of the EMR section, was ablated with a laser (400 Hz).

### 6. Stitching & registration of IMC images to IF images

We developed a two-step approach for the stitching and registering of dual-modality imaging data. For each small FOV of IMC, the normalized cross-correlation of the IMC DNA image and IF DAPI is first calculated. The peak of the cross-correlation matrix identifies the location where the small FOV IMC image is best correlated with the subregion of WSI IF. Then, feature-based image registration is performed by identifying matched features from each imaging modality^26^. Each FOV image can be mapped onto WSI IF and stitched together. High-resolution of the segmentation mask from IF image is also registered by matching the resolution of the IMC image.

### 7. Visualization

The registered IF images (n=3) and IMC images (n=36) were simultaneously visualized by open-source visualization software, Napari (https://github.com/napari/napari, Supplementary Figure 2)

### 8. Single-cell segmentation

We performed cell and nuclear segmentation using Mesmer^16^ using a nuclear marker (DAPI for IF and DNA for IMC) and an epithelial membrane marker (E-Cadherin, Figure 4).

### 9. Feature extraction

We used a cell segmentation mask made from IF images to obtain the mean intensity of high-dimensional IMC image sets (Figure 4). By applying the newly selected region of interest corresponding to each disease status (Figure 3A, B), we extracted the mean intensity of 36 IMC antibodies using a single cell mask generated from high-resolution IF images (Figure 4E). Feature extraction was performed using the Skimage python library (https://scikit-image.org). More cells were segmented using IF masks (DAPI, E-Cadherin, total: 199,425 cells) than by using IMC masks (DNA, E-Cadherin, total: 120,643 cells)

### 10. Unsupervised clustering

We performed cell phenotyping from the single-cell data using a GPU-boosted implementation of PhenoGraph^7^ (Figure 7A).

## Supplemental information titles and legends

**Supplementary Figure 1.**
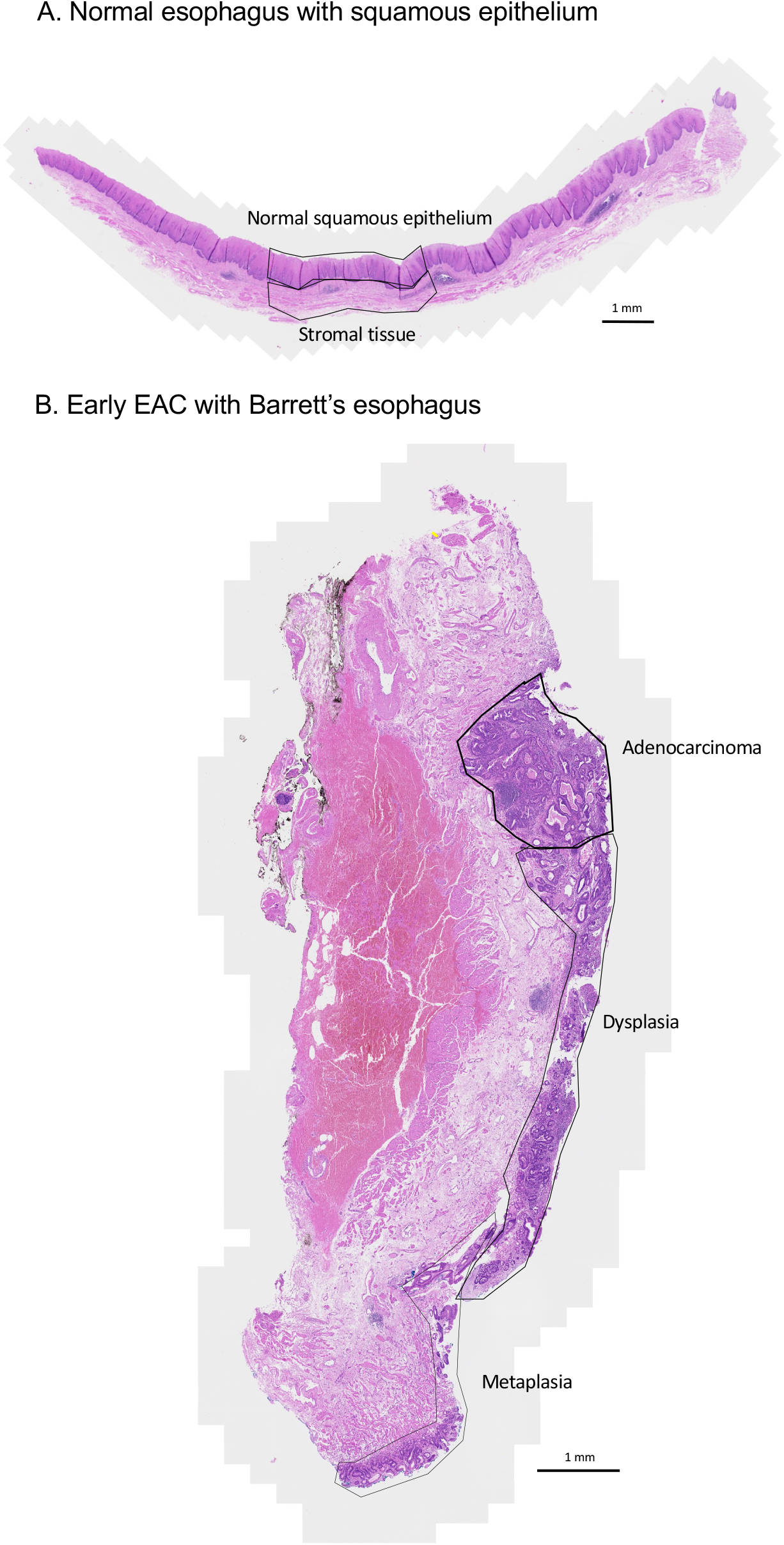
Representative H&E image of EMR section. A. Normal esophagus with squamous epithelium. B. Early EAC arising from Barrett’s esophagus

**Supplementary Figure 2.**
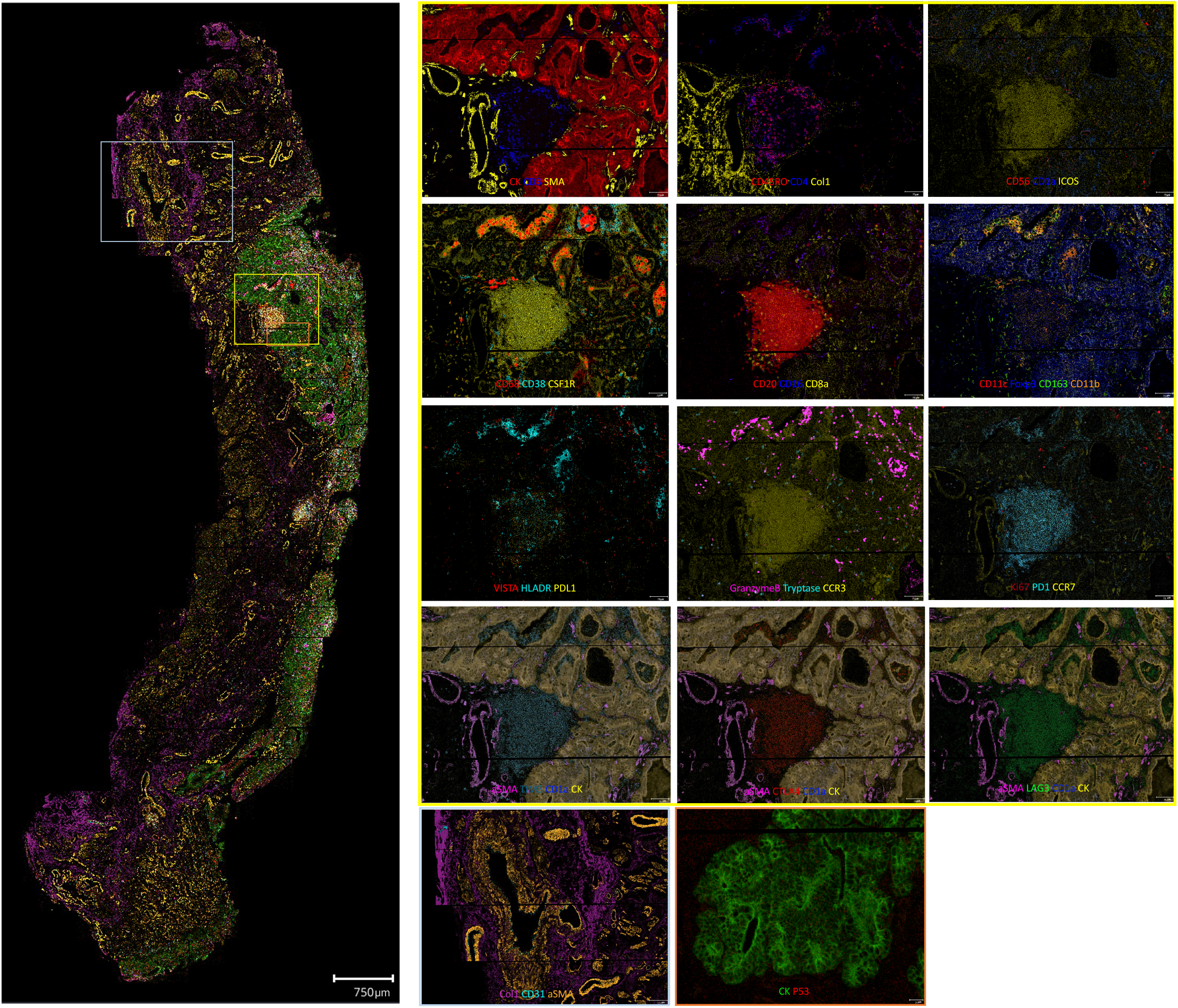
Visualization for 36 antibodies of IMC markers

**Supplementary Figure 3.**
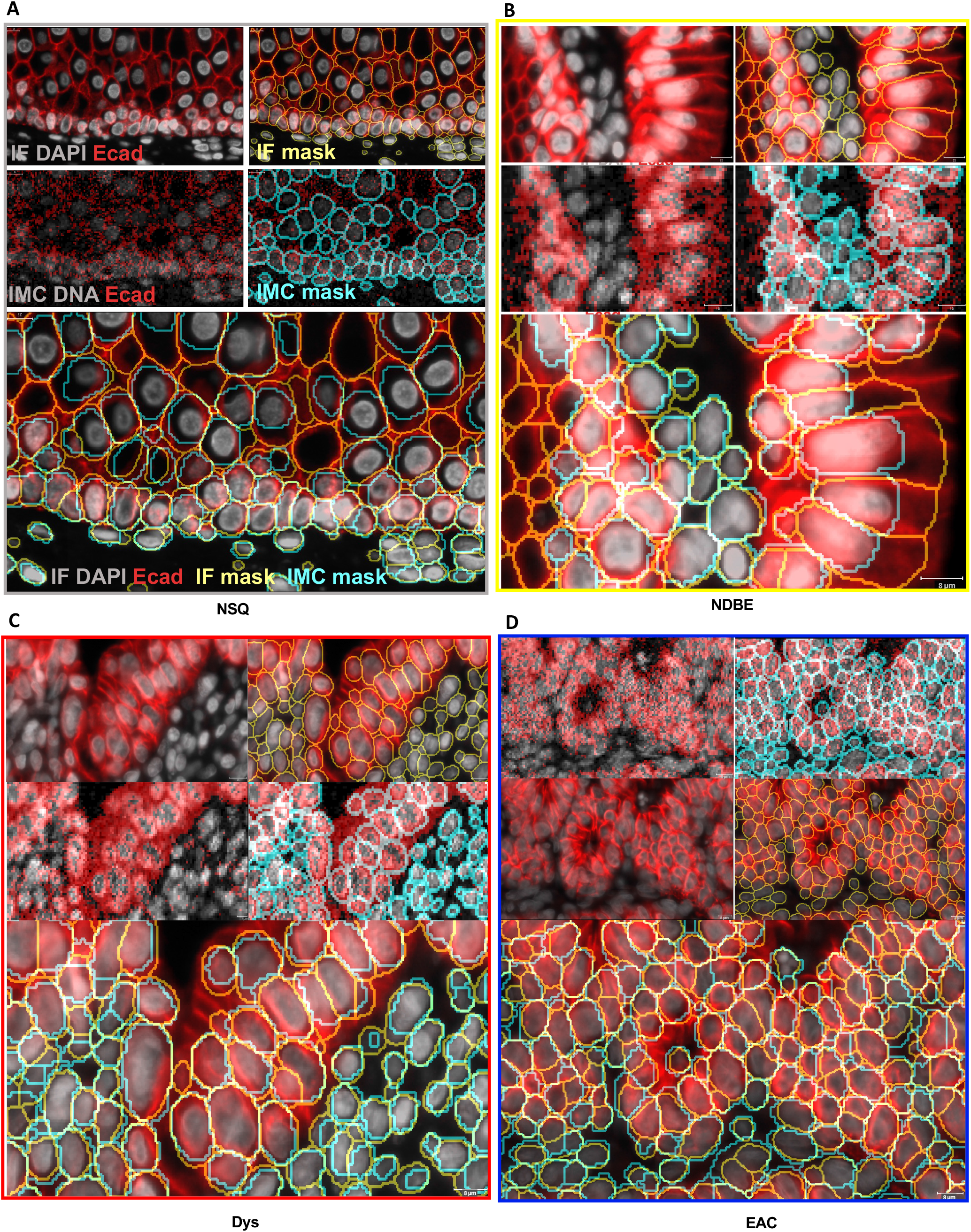
Segmented cell mask comparison between IF-based and IMC-based according to disease progression A. Normal squamous epithelium (NSQ), B. Nondysplastic Barrett esophagus (NDBE), C. Dysplasia (Dys), D, esophageal adenocarcinoma (EAC)

**Supplementary Figure 4.**
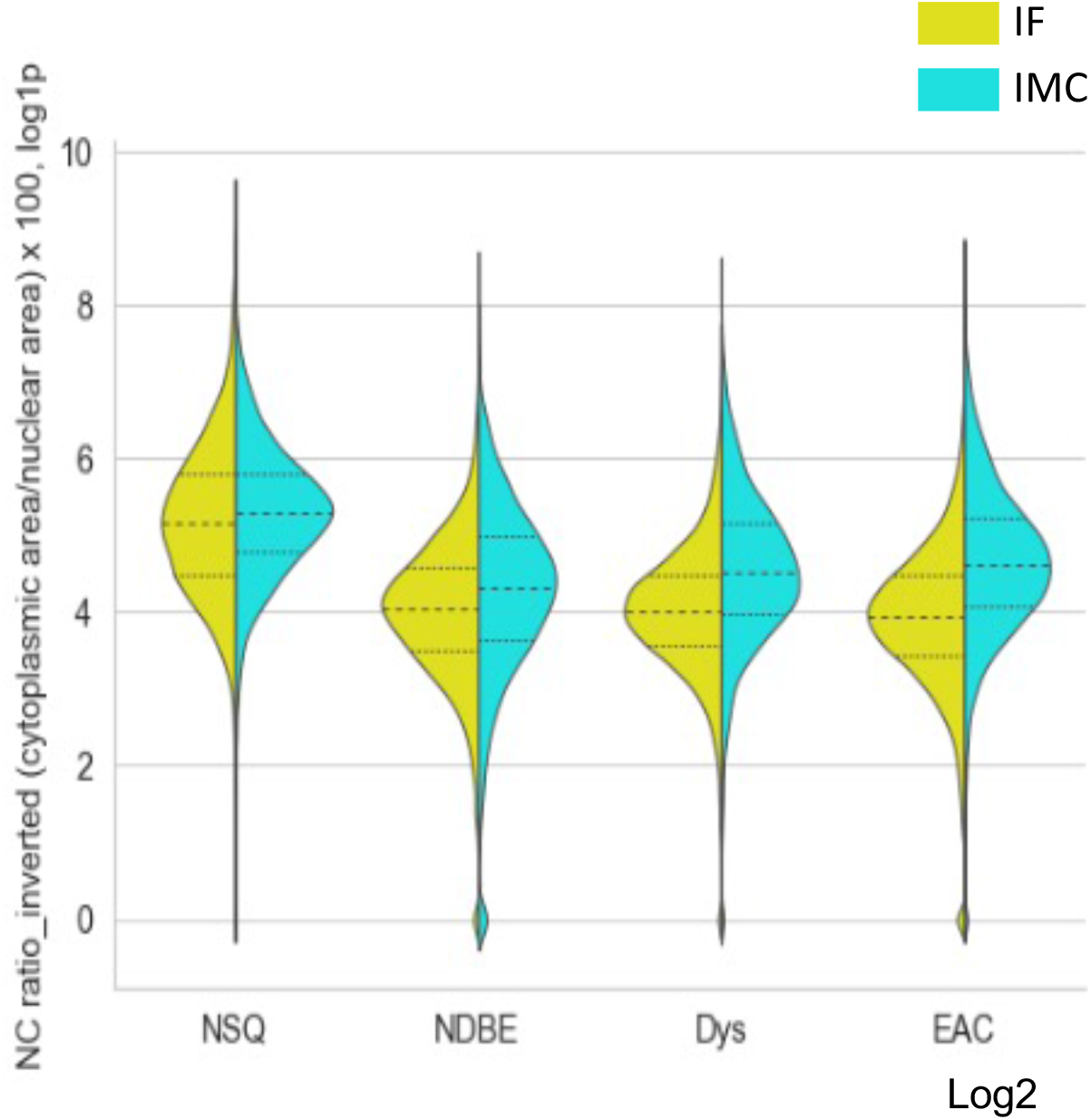
Cytoplasmic/Nuclear ratio (inverted NC ratio) across disease progression in all regions

**Supplementary Figure 5.**
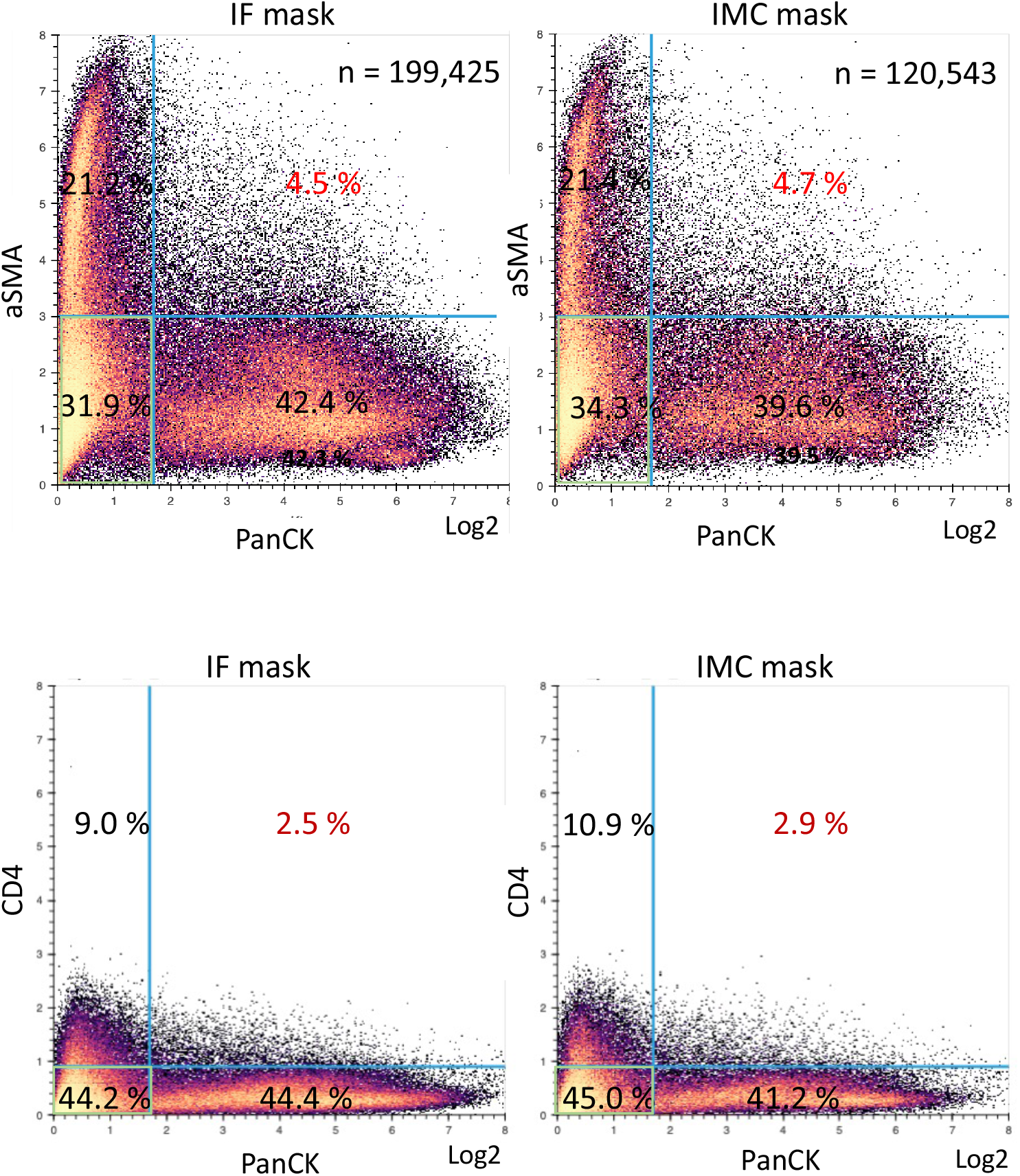
Biaxial scatter plot of mean intensity

**Supplementary Figure 6.**
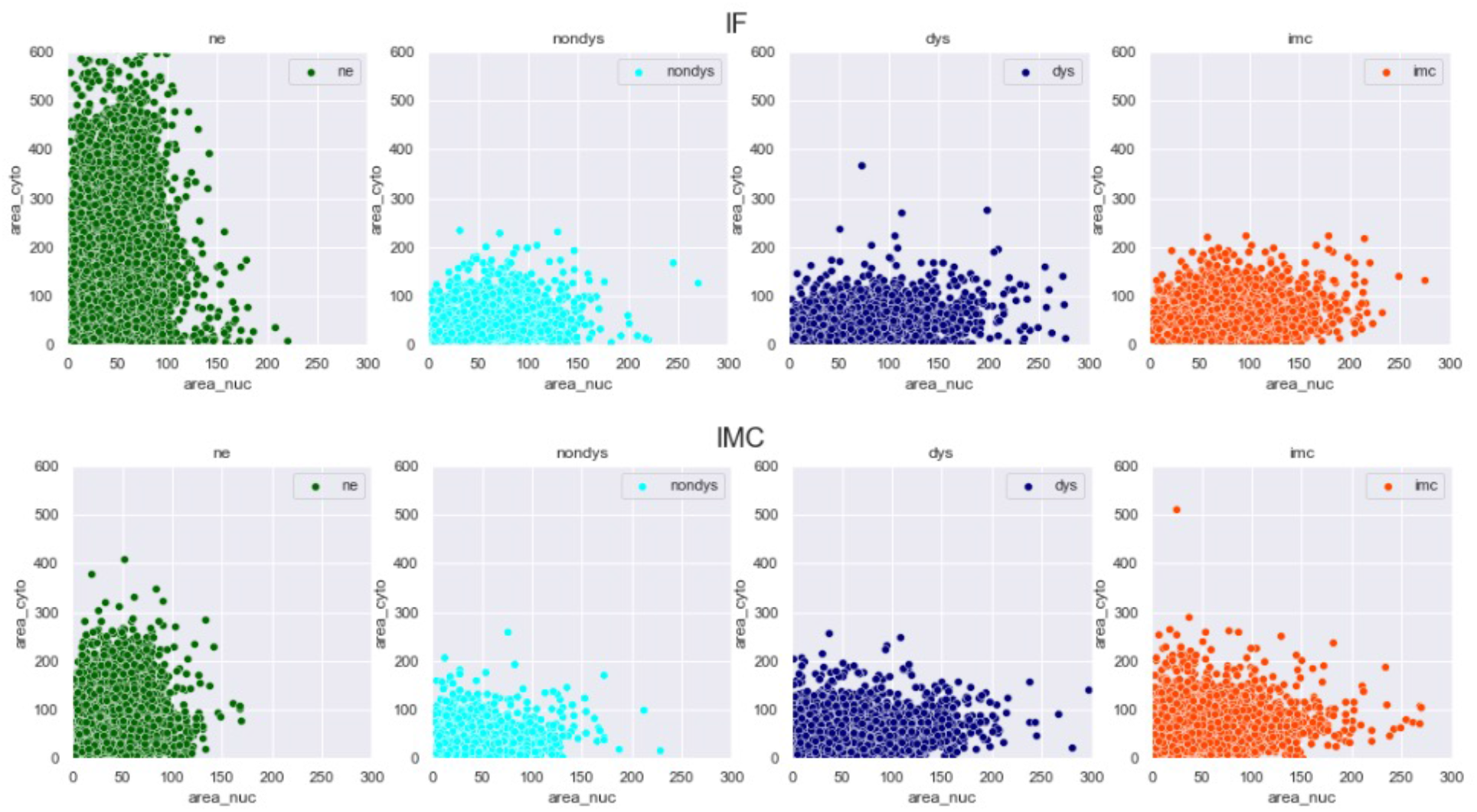
Nuclear area vs. cytoplasmic area according to the disease status

## Figures, supplemental information, and the key resources

### KEY RESOURCES TABLE

**Supplementary Table 1.** 36 metal-tagged antibodies for IMC.

| Class | No. | Metal |  | Clone | Type | Cat No. | Antibody | Name | Comments |
| --- | --- | --- | --- | --- | --- | --- | --- | --- | --- |
| 1. Epithelial cell | 9 | Nd | 148 | C11 | Human | 3148020D | Pan-Keratin | panK | Epithelial cells |
|  | 18 | Gd | 158 | 24E10 | Cross | 3158029D | E-Cadherin | E-Cad | Epithelial cells |
| 3. Stromal - vascular cell | 2 | Pr | 141 | 1A4 | Human | 3141017D | Alpha-SMA | aSMA | Vascular, fibroblast, smooth muscle |
|  | 29 | Tm | 169 | Polyclonal | Human | 3169023D | Col 1 | Col1 | Extracellular matrix |
|  | 3 | Nd | 142 | EPR3094 | Rabbit IgG | ab207090 | CD31 | CD31 | endothelial cells |
| 2. P53 mutation proliferating tumor cells | 4 | Nd | 143 | DO-7 | Human | 3143026D | p53 | p53 | p53 protein accumulation |
|  | 28 | Er | 168 | B56 | Cross | 3168022D | Ki-67 | Ki67 | Proliferating cells |
| 4. Lymphoid Lineage (T cell) | 33 | Yb | 173 | UCHL1 | Cross | 3173016D | CD45RO | CD45RO | Memory T cells |
|  | 30 | Er | 170 | Polyclonal, C-Terminal | Human | 3170019D | CD3 | CD3 | pan T cells |
|  | 17 | Gd | 156 | EPR6855 | Human | 3156033D | CD4 | CD4 | Helper T cells |
|  | 10 | Sm | 149 | Y31 | Rabbit IgG | ab228341 | CCR3 | CCR3 | Helper T cell : Th9 cells |
|  | 32 | Yb | 172 | Y59 | Rabbit IgG | ab221209 | CCR7 | CCR7 | Naïve T cell/ Memory(high) T cells |
|  | 35 | Lu | 175 | EP2567Y | Rabbit IgG | ab215981 | CD56 | CD56 | NK cell, NK/T cells |
| 5. T cell,<br>Functioning<br>Cell | 22 | Dy | 162 | C8/144B | Human | 3162034D | CD8a | CD8a | Cytotoxic T cells |
|  | 27 | Er | 167 | EPR20129-217 | Human | 3167021D | Granzyme<br>B | GranB | Activated T<br>cells/NK cells |
| 6. Lymphoid<br>cells<br>- regulatory<br>or exhausted | 20 | Gd | 160 | EPR20560 | Rabbit<br>IgG | ab225577 | ICOS | ICOS | Helper T cell :<br>TFH cell s |
|  | 34 | Yb | 174 | EPR22241 | Rabbit<br>IgG | ab242080 | TIM 3 | TIM3 | Exhausted T<br>cells |
|  | 12 | Eu | 151 | EPR20261 | Rabbit<br>IgG | ab227579 | LAG-3 | LAG3 | Exhausted T<br>cells |
|  | 26 | Er | 166 | CAL49 | Rabbit<br>IgG | ab251599 | CTLA4 | CTLA4 | Regulatory T<br>cells |
|  | 16 | Gd | 155 | 236A/E7 | Human | 3155016D | FoxP3 | Foxp3 | Regulatory T<br>cells |
|  | 24 | Dy | 164 | D4W2J | Rabbit<br>IgG | 86163BF | PD-1 | PD1 | TFH cell,<br>Exhausted T<br>cells |
| 7. Lymphoid<br>cells<br>- B<br>cell/plasma<br>cell lineage | 21 | Dy | 161 | H1 | Human | 3161029D | CD20 | CD20 | B cells |
|  | 6 | Nd | 145 | EPR4106 | Rabbit<br>IgG | ab176886 | CD38 | CD38 | NK cells,<br>monocytes,<br>activated B<br>cells/T cells |
| 8. Myeloid<br>- Neutrophil,<br>mast cell | 5 | Nd | 144 | EPR1344 | Rabbit<br>IgG | ab209970 | CD11b | CD11b | Pan-myeloid<br>cells |
|  | 7 | Nd | 146 | EPR16784 | Rabbit<br>IgG | ab215977 | CD16 | CD16 | Neutrophil |
|  | 31 | Yb | 171 | EPR9522 | Rabbit<br>IgG | ab271916 | Mast Cell<br>Tryptase | Tryptas | Myeloid- mast<br>cells |
| 9. Myeloid<br>-<br>macrophage | 36 | Yb | 176 | EP1347Y | Rabbit<br>IgG | ab216655 | CD11c | CD11c | Dendritic cells |
|  | 1 | Qdot<br>800 | 112/114 | EPR3653 | Rabbit<br>IgG | ab226121 | CD14 | CD14 | Monocytes |
|  | 23 | Dy | 163 | EPR3692 | Rabbit<br>IgG | ab215985 | HLA-DR | HLA-DR | macrophage-APC (MHC II) |
|  | 19 | Tb | 159 | KP1 | Human | 3159035D | CD68 | CD68 | Macrophages |
|  | 8 | Sm | 147 | Edhu-1 | Human | 3147021D | CD163 | CD163 | Macrophages-M2 |
|  | 14 | Eu | 153 | EPR22489-7 | Rabbit IgG | ab254471 | Mannose Receptor | CD206 | Macrophage |
|  | 13 | Eu | 152 | O10 | Mouse IgG1 | ab212980 | CD1a | CD1a | Langerhans cells |
|  | 15 | Sm | 154 | SP211 | Rabbit IgG | ab240265 | CSF-1-R | CSF1R | CSF1R macrophages |
| 10. T cell inhibition | 11 | Nd | 150 | E1L3N | Human | 3150031D | PD-L1 | PDL1 | Immune checkpoint Ligand, Inhibit T cells |
|  | 25 | Ho | 165 | EPR21050 | Rabbit IgG | ab238886 | VISTA | VISTA | Inhibit T cells/ inhibit myeloid cell |

## Notes

### Competing Interest Statement

The authors have declared no competing interest.

